# CompareM2 is a genomes-to-report pipeline for comparing microbial genomes

**DOI:** 10.1101/2024.07.12.603264

**Authors:** Carl M. Kobel, Velma T. E. Aho, Ove Øyås, Niels Nørskov-Lauritsen, Ben J. Woodcroft, Phillip B. Pope

**Affiliations:** Faculty of Biosciences, Norwegian University of Life Sciences, Ås, Norway; Clinical Institute, University of Southern Denmark, Odense, Denmark; Centre for Microbiome Research, School of Biomedical Sciences, Queensland University of Technology (QUT), Translational Research Institute, Woolloongabba, Australia; Faculty of Chemistry, Biotechnology and Food Science, Norwegian University of Life Sciences, Ås, Norway

**Keywords:** microbiology, bacteria, archaea, genomics, metagenome assembled genomes, bioinformatics, pipeline, workflow, genomic annotation, phylogenetics, parallel computing

## Abstract

Here, we present CompareM2, a genomes-to-report pipeline for comparative analysis of bacterial and archaeal genomes derived from isolates and metagenomic assemblies. CompareM2 is easy to install and operate, and integrates community-adopted tools to perform genome quality control and annotation, taxonomic and functional predictions, as well as comparative analyses of core- and pan-genome partitions and phylogenetic relations. The central results generated via the CompareM2 workflow are emphasized in a portable dynamic report document. CompareM2 is free software and welcomes modifications and pull requests from the community on its Git repository at https://github.com/cmkobel/comparem2.

## Background

Costs are decreasing both for sequencing of microbial genomes and complex microbiomes and for the computational resources necessary to analyze generated reads. This has led to an exponential growth in the number of available genomes and metagenome-assembled genomes (MAGs). Despite this growth, there are limits on the accessibility of software that can analyze the evolutionary relationships and functional characteristics of microbial genomes in order to assess variation of both known and unknown species. Much of the software commonly used to analyze prokaryotic genomes has a high user entry level, requiring advanced skills for complicated installation procedures, debugging dependency issues, and circumventing operating system-specific limitations. This results in a disproportionate amount of time being spent by researchers on setup and technical preparations needed to analyze the sequenced genomic reads rather than biologically relevant analysis of scientific data. These factors define the backdrop that has motivated the conceptualization, development, and application of the CompareM2 genomes-to-report pipeline, which is designed to be an easy-to-install, easy-to-use bioinformatic pipeline that makes extensive analysis and comparison of microbial genomes straightforward.

We compared CompareM2 to several other pipelines that are designed for overlapping use cases: Nullarbor^1^, Tormes^2^ (stylized TORMES) and Bactopia^3^ (**Table 1**). Nullarbor and Tormes do assembly and comparison and have a focus on antimicrobial resistance, spread of pathogens, and core genomes relevant for analyzing individual species. They both produce a report document that is similar to what CompareM2 produces. Bactopia does both assembly and comparative analyses, but while it does some comparative analyses in conjunction with assembly, the user must launch individual predefined workflows included in the Bactopia Tools extension to compare between the samples. Bactopia does not have a parallel scheduler for running these comparative tools. While it does not produce a report document, it does have more overlapping tools with CompareM2 when considering the Bactopia Tools extension. Neither Tormes nor Bactopia is designed for analyzing archaea, although many of the tools integrated in these pipelines are applicable for archaeal genomes when care is taken, e.g. core/pan genome reconstruction and phylogenetic analysis, etc.

**Table 1:**
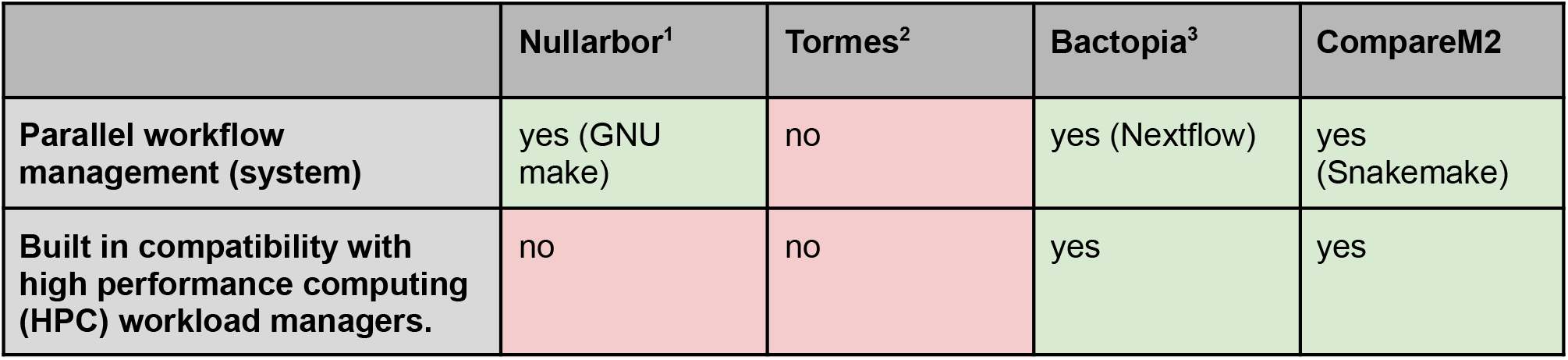
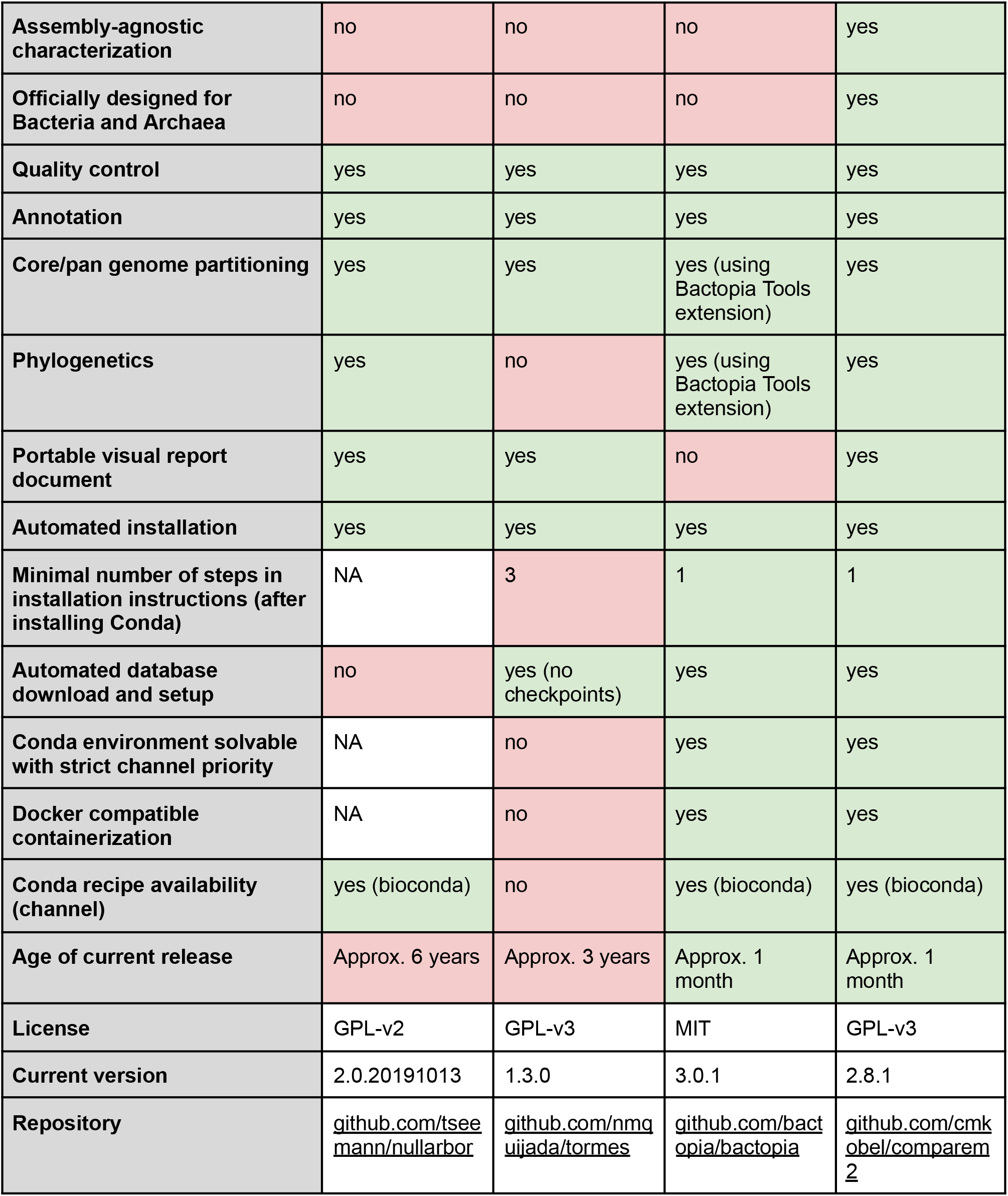
Qualitative comparison of Nullarbor, Tormes, Bactopia and CompareM2.

Furthermore, there is a lack of tools to analyze archaea which means that in many cases, researchers may opt to use non-archaeal tools for analysis of these. For this reason we have opted to compare them to CompareM2, which is designed to analyze both bacteria and archaea. Tormes has a sequential architecture, which means that it runs one sample at a time and one tool at a time. This is in contrast to CompareM2 and Bactopia, which have a parallel job scheduler where several samples and tools can be run at the same time. CompareM2 inherits this property from Snakemake, on which it is built. Bactopia on the other hand is built on the Nextflow workflow system which in many cases is comparable to Snakemake. Central processing units (CPUs) of computers, whether in laptops, workstations, or HPCs, are seeing an increasing number of physical cores. To take advantage of this, it is necessary for software to have a parallel architecture that can utilize the full potential of the processing resources available. This is especially important on HPCs, where many independent compute nodes can run parallel jobs in a scalable manner.

Another bottleneck in bioinformatics is the interpretation of large output files and visualization of data in an informative manner. CompareM2 produces a graphical report that contains the most important curated results from each of the analyses carried out on the user-specified set of query genomes. This report contains text and figures that explain the significance of the results, which makes it easy to interpret for users with a non-bioinformatics background.

While CompareM2 can be used to compare prokaryotic isolate genomes, it also contains tools to analyze bins or MAGs from the sequencing of large microbial communities. The genome is the foundation of any multi-omics study, and such a resource of annotated genomes can be readily integrated into subsequent multi-omics analyses. For example, metaproteomic searches require a highly specific and well-annotated genome database to match MS/MS spectral data^4,5^.

## Results

CompareM2 congregates the most commonly used and community-tested tools to perform prokaryotic genome quality control, gene calling, functional annotation, phylogenetic analysis, and comparison of genomes across the core-pan spectrum. A major priority of CompareM2 is the ease of installation and use, which is achieved by containerizing all bundled software packages and automatizing the download and setup of databases. The choice of genomes to input can be any set where there is a comparable feature either within or between species. The number is limited by the computational resources but the dynamic report is designed for comparing hundreds of genomes.

### Software design

CompareM2 is written as a command line program that the user calls with the input genomes that they wish to analyze. It has a text interface where the user can define optional parameters and a single executable that takes care of the overall procedure: First, it checks for presence of the Apptainer runtime, and defines reasonable defaults for database directories and configuration files, in case the user has not specified these manually as environment variables. There also is a “passthrough arguments” feature that makes it possible to address any command line argument to any rule in the workflow. (further details in documentation https://comparem2.readthedocs.io/en/latest/). One example of a setting that can be defined via the configuration file is whether to optionally submit jobs through a workload manager like Slurm, PBS, etc., typically used on high-performance computing clusters (HPCs). Next, the executable dispatches the main Snakemake pipeline that runs all genomic analyses.

This main pipeline automatically installs all necessary software environments and automatically downloads necessary databases, depending on which analyses the user has selected to run. Finally, it dispatches rendering of the dynamic report which contains the results of the main pipeline. This report is dynamic in the sense that it only includes the results which are present, which means that it can be rendered independently of which analyses the user has selected to compute.

Overall, CompareM2 is designed in such a way that the user can install the complete software in a single step. Similarly, running all analyses on a set of microbial genomes (bacterial and archaeal) can be launched in a single action, and the curated results can be studied in the dynamically rendered report. The machine requirements are a Linux-compatible OS with a Conda-compatible package manager, e.g., Miniforge, Mamba or Miniconda. There is nothing standing in the way of running CompareM2 on other operating systems, but many of the included bioinformatic tools for genomic analysis are mostly compatible with Linux-like x64-based systems. For a technical description of how CompareM2 is implemented, please see the Methods section.

For demo reports, please seehttps://comparem2.readthedocs.io/en/latest/30%20what%20analyses%20does%20it%20do/#rendered-report.

## Benchmarking

Initially, we wanted to compare CompareM2 to Nullarbor^1^, Tormes^2^ and Bactopia^3^. As none of these tools support the external long-reads based assembly, binning and dereplication pipeline where our MAGs were sourced from, we inputted the finished MAGs as is into these tools. Unfortunately this was not possible for Nullarbor, as it is not able to run without reads^6^. Nonetheless, we have included Nullarbor in **Table 1** for the purpose of a qualitative comparison.

We compared the running times of CompareM2, Tormes, and Bactopia when scaling up the number of input MAGs to analyze on a single workstation. We considered two different genera: *Methanobrevibacter*, which are archaea from the class Methanobacteria, and *Prevotella*, which are Gram-negative bacteria from the class Bacteroidia. Our MAGs have an average genome size of 2.19 Mb for *Methanobrevibacter* and 3.07 Mb for *Prevotella*.

Species prediction and genome sizes are measured on the analyzed MAGs with GTDB-Tk^7^ and assembly-stats^8^ using CompareM2 itself.

Although Bactopia, Tormes, and CompareM2 are designed for overlapping use cases, they are still very different, because they implement different kinds of analyses. In order to make them as comparable as possible, we ran only the analyses with pairwise overlap between CompareM2 and each of the two other tools. This was done using CompareM2’s “until” parameter to specify exactly which rules to run. CompareM2 in “Bactopia mode” includes rules sequence_lengths, assembly_stats, prokka, abricate, and mlst, whereas CompareM2 in “Tormes mode” includes rules prokka, abricate, assembly_stats, mlst, panaroo, and gtdbtk.

We analyzed the running time of Bactopia, Tormes and CompareM2 when increasing the input size (number of input genomes) (**Fig. 1**). The running times of all tools were approximately linear functions of the input size (time = input size × slope + constant) but with big differences between pipelines in the slope. There are hints of an exponential component in the scaling of running time for Tormes and CompareM2 in Tormes mode, likely because these pipelines construct core genomes, which is a computationally expensive problem where all genes, in the case of Panaroo and Roary, are compared in a pairwise manner^9,10^. The running time per genome was generally higher for *Prevotella* than *Methanobrevibacter*, which is expected since *Prevotella* has a slightly larger average genome size, meaning that the total number of genes to be processed is larger.

**Fig. 1:**
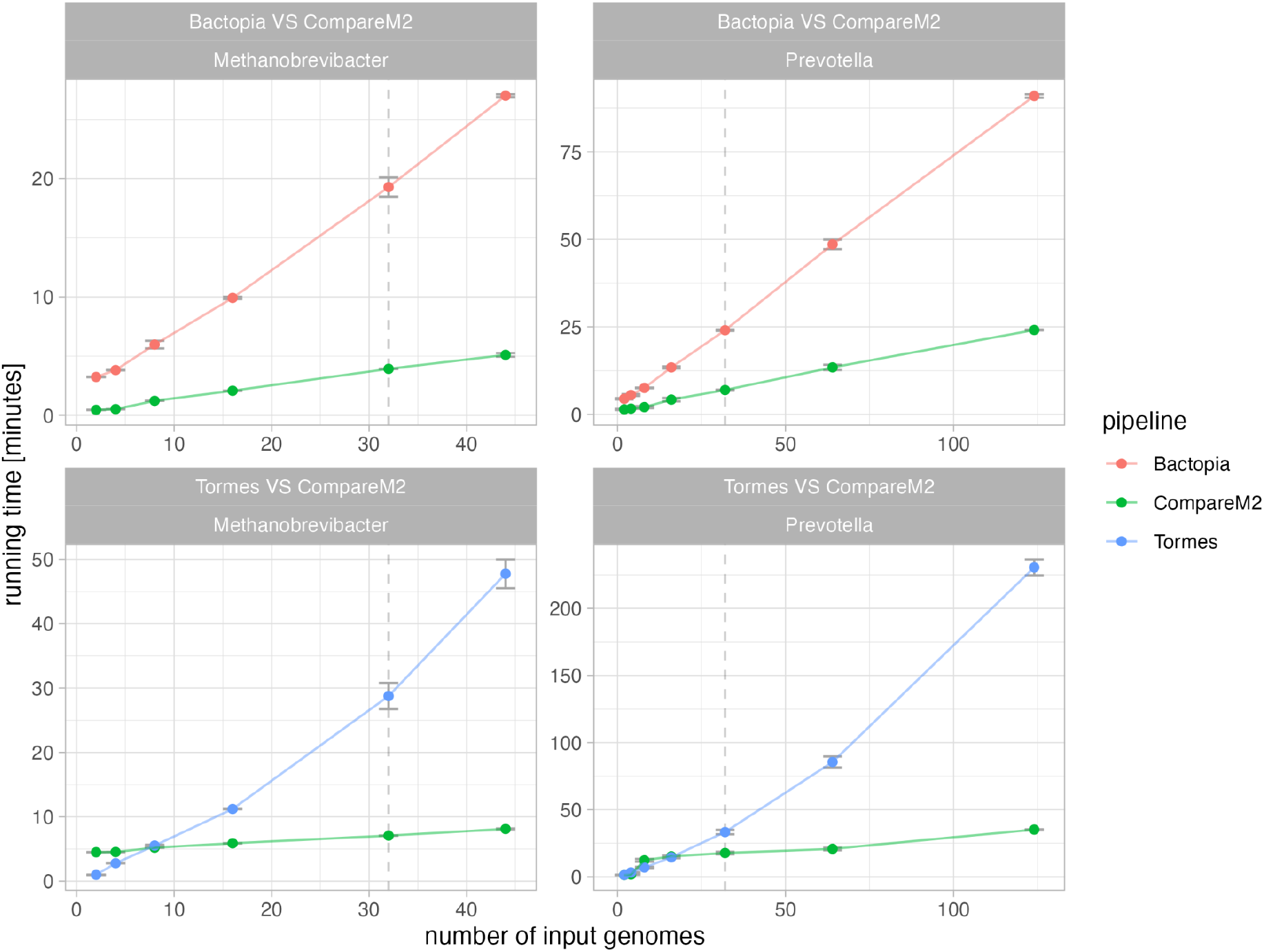
Wall running time analysis comparing CompareM2 to Bactopia and Tormes. In each comparison, CompareM2 was run in a mode where it creates a comparable set of results to the pipeline it is compared to. All analyses ran with 3 replicates, the error bars show means ± the standard deviation of these replicates. A vertical dashed line highlights input size = 32 which is equal to the number of cores used in each benchmark.

For running time per number of input genomes, CompareM2 outperformed both Tormes and Bactopia significantly. When analyzing 44 *Methanobrevibacter* MAGs, CompareM2 is 4.1 times faster than Bactopia and 7.2 times faster than Tormes. For 124 *Prevotella MAGs*, CompareM is 3.1 and 7.8 times faster than Bactopia and Tormes, respectively (**Table 2**).

**Table 2:** Wall running time in seconds for analyzing 44 Methanobrevibacter MAGs or 124 Prevotella MAGs. “Factor” denotes how many times slower each tool is compared to the fastest. The fastest tool is marked with an underline (in both cases CompareM2). All numbers are means of three replicates.

|  | 44 *Methanobrevibacter* genomes |  | 124 *Prevotella* genomes |  |
| --- | --- | --- | --- | --- |
|  | minutes | factor | minutes | factor |
| **<u>CompareM2</u>** | <u>6.6</u> | <u>1.0</u> | <u>29.7</u> | <u>1.0</u> |
| **Bactopia** | 27.0 | 4.1 | 91.0 | 3.1 |
| **Tormes** | 47.7 | 7.2 | 230.5 | 7.8 |

## Discussion

CompareM2 is significantly faster than both Tormes and Bactopia as its running time scales much better with increasing input size. Notably, running time scaled approximately linearly with a small slope even when increasing the number of input genomes well beyond the number of available cores on the machine. The running time of each pipeline comes down to the time it takes to run each included tool on each sample, so differences between pipelines in terms of running time are determined by how they allocate resources and schedule jobs efficiently in parallel.

The speed of Bactopia is strongly affected by its reads-based approach: If reads are not input by the user – which was not possible in this case because we compared genomes that were assembled using a different pipeline – Bactopia creates artificial reads with ART^11^. This is done in order for Bactopia to be able to compare genomes without reads to genomes with reads. CompareM2 on the other hand is designed to compare genomes without reads and thus does not have to spend computing resources on producing these artificial reads. It should be noted that if the user runs more comparative analyses using the Bactopia Tools extensions, the scalability will be worse since the Bactopia platform does not offer to schedule running several tools in parallel. While Tormes does not suffer from producing artificial reads, it does fall short on not having a parallel workflow management system. As it runs all samples sequentially, running each tool at a time, it is not competitive on HPCs or multi-core CPUs. Generally, the running time standard deviations are negligible because the relative time differences are large. The running time was computed on a 64-core workstation (see Methods - Benchmarking). We ran the analysis by allocating 32 cores on this machine. By running with a lower number of cores than the machine has, we lower the chances that any other component than the CPU is the main bottleneck for computation.

Since both Tormes and Bactopia are designed for different use cases, they might not represent the perfect contenders for a comparison with CompareM2. Nonetheless, to our knowledge, they are the most comparable pipelines that exist today. In the case of Tormes, the comparison highlights the benefit of having a parallel rather than sequential job scheduling setup. In the case of Bactopia, it shows that other pipelines can approach the scalability of CompareM2 but also that having a reads-based approach is not competitive and that comparative analyses can be more integrated into the main pipeline. Also, we want to highlight that Bactopia and Tormes are not the only tools relevant for comparison. As CompareM2 sports many tools for advanced annotation, it also overlaps in use case with more annotation-focused pipelines like DRAM^12^.

What is characteristic about CompareM2, is that it is assembly-agnostic: It works strictly downstream of assembling and binning. It is a general-purpose pipeline that doesn’t discriminate between genomes based on how they were assembled. Many other tools also include all the steps necessary to turn raw reads into genome representatives and then do varying degrees of biological characterization of these, but raw read-dependent tools were deliberately left out of CompareM2. This is because read mapping, assembling, or even binning are highly dependent on the sequencing technology used and require a highly specialized pipeline for each technological use case. Next-generation sequencing has matured, and many competitive sequencing platforms exist (sequencing-by-synthesis, single molecule sequencing, etc.). Thus, designing a toolbox that can compare genomes is a very different discipline from designing a toolbox that can assemble these genomes in the first place. Hard-linking two such pipelines together raises the concern that one part will not fit a specific use case. CompareM2 takes a different approach which is to offer a platform where you can compare your genomes regardless of how they were assembled.

## Conclusions

CompareM2 offers an easy-to-install, user-friendly, and efficient genome annotation pipeline. It can be launched using a single command and is scalable to a range of projects, from the annotation of single genomes to comparisons across complex inventories. By using widely adopted genome tools, CompareM2 performs key annotation steps including genome quality control, predicted biological gene function, and taxonomic assignment. In addition, comparative analyses like computation of core- and pan-genomes or phylogenetic relations can be executed. We expect that CompareM2 will support the productivity of genome researchers by simplifying and expediting the annotation and comparison of genome-centric data. Further development of CompareM2 will continue with its ongoing adaptation to the community consensus of microbial ecologists. Through benchmarking, we have shown that CompareM2 is highly scalable, allowing analysis of large numbers of input genomes thanks to its underlying parallel job scheduling provided by Snakemake. Via CompareM2 we seek to accelerate and democratize the analysis of genomic assemblies for anyone who has computational resources available—be that on HPCs, a workstation, or even a laptop.

## Methods

### Implementation of CompareM2

#### Snakemake

*This section sketches the technical implementation of CompareM2 using Snakemake. For more details on usage and modification of running parameters, please consult the documentation at: https://comparem2.readthedocs.io*.

CompareM2 is built on top of Snakemake^13^. This means that much of the functionality in it is inherited by solutions within the Snakemake framework. A major upside of this is that CompareM2 can make use of Snakemake’s extensive support for parallel job scheduling and support for running on workload managers used on HPCs. CompareM2 has been tested on Slurm^14^ and PBS^15^ workload managers. The CompareM2 executable, which is available on Bioconda, is run by the user and does some basic housekeeping: It sets up the necessary environment variables like the base directory of the code, the Snakemake profile that defines whether the jobs are being submitted to an HPC workload manager and whether to use Conda or Docker/Apptainer as well as checking the database paths which are used by some of the tools. Then, the main Snakemake workflow is launched on the input genomes. By default CompareM2 accepts all fasta-formatted files available in the current working directory, but a specific path of “input_genomes” can also be set. In the case of analyzing large numbers of genomes, a “fofn” (file of file names) can also be used to define the input genomes. Snakemake allows for running only specific wanted parts of the rule graph (**Fig. 2**) and common parameters can be customized by the user. Because command line arguments are passed on from the CompareM2 executable to the main Snakemake workflow, the user has full access to control the main Snakemake workflow underneath.

**Fig. 2:**
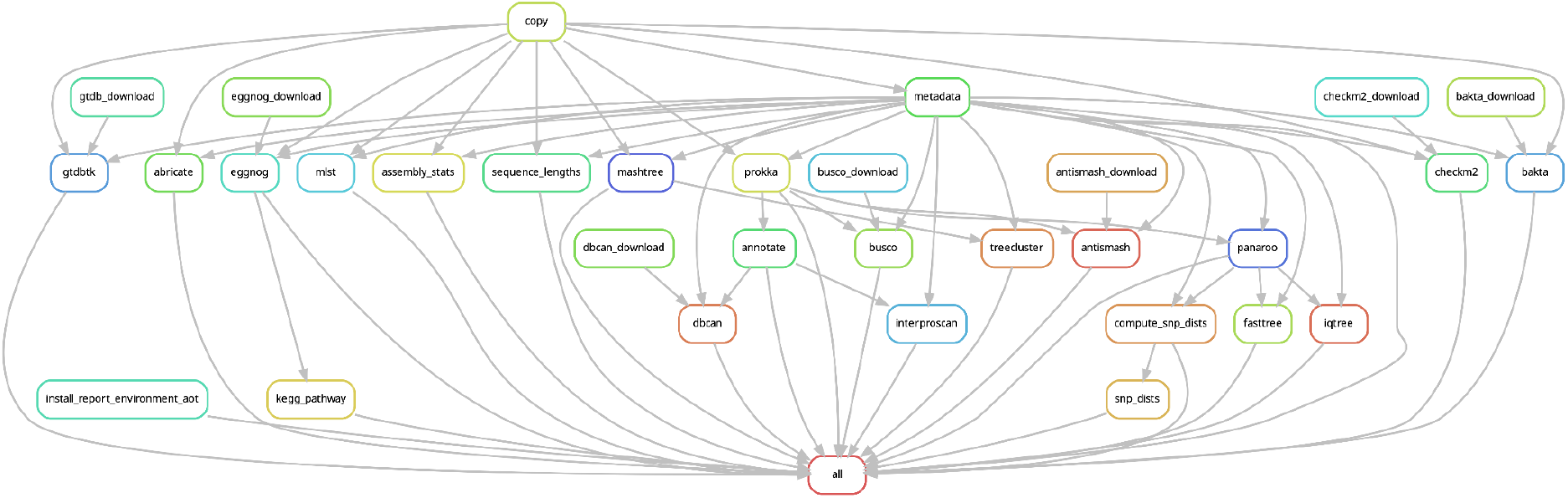
The underlying Snakemake workflow of CompareM2. A directed acyclic graph (DAG) that shows the order in which to run rules that are dependent on each other. Each box represents a Snakemake rule, which is a code template that can be used to process a sample or a batch of samples. For instance, rule “prokka” is run on each input assembly, and then rule “panaroo” can analyze all samples using this output. The start and end points for the complete pipeline are represented by “copy” and “all”, respectively.

Regardless of whether the user uses the precompiled Docker image that contains environments for the included tools, the snakefile rules that govern how the specific tools are called can still be fully modified.

When the main workflow is completed, the CompareM2 executable checks that relevant outputs exist, and in that case it calls the “dynamic report” sub pipeline. This pipeline is only in charge of producing the portable graphical dynamic report document that contains the main results of the main pipeline. When this report is rendered, the return code of the main pipeline is returned by the CompareM2 executable. One of the main features of this dynamic report is that it can produce a report from possibly incomplete results from the main pipeline. In many cases, a job will fail; for instance an advanced annotator like Antismash^16^ will return a non-zero exit code and not produce any output files if no genes are annotated in a genome. When this is the case, it is preferable that the rest of the main pipeline can continue and that the dynamic report pipeline can be run on the remaining results, showing the missing results. In a pipeline with this many moving parts, it is crucial that unaffected tools can continue running if something breaks down. This graphical report is based on the R Markdown^17^ document rendering framework. Statistics and plots are generated with Tidyverse^18^.

Launching the pipeline is a single command and the user is only required to have minimal experience with the command line interface for moving fasta genome files in and out of directories. The minimal system requirements are Linux with a Conda-compatible package manager. It is recommended that the system has Apptainer installed such that a Docker image containing all necessary binaries can be automatically downloaded and installed.

#### Docker and downloads

As CompareM2 uses many independent tools for analysis, it is not feasible to have all binaries in the same environment. The requirements for many different versions of the same dependencies would quickly lead to dependency hell^19^. Instead, each tool is installed into its own isolated environment. For a developmental installation of CompareM2, these isolated Conda environments are automatically installed using the “use-conda” functionality from Snakemake. In any other case when a user installs CompareM2 and has Apptainer (a high-performance Docker-compatible runtime^20^), the “use-apptainer” system is activated and a pre-compiled Docker^21^ image is automatically downloaded and used instead. It contains a precompiled distribution of all Conda environments. Using this image is optional but highly recommended as it avoids potential dependency issues for the user to deal with during installation. Six of the bundled tools (Bakta, Busco, CheckM2, Dbcan, Eggnog, GTDB-Tk) need databases to run. These databases are automatically downloaded by individual rules in the Snakemake rule graph. CompareM2 only downloads these databases the first time a tool is used.

#### Tools included

Many tools are included in CompareM2. Here we list which they are, what they do and define the conditions in which they run.

The first step of the pipeline is to run all genomes through any2fasta^22^ which acts as input validation and converts the input genome queries into a homogenized fasta format with a uniform character set.

Quality control is performed by assembly-stats^8^ and seqkit^23^ which both compute various basic genome statistics like genome length, count and lengths of contigs, N50, GC, etc.

Busco^24^ and CheckM2^25^ are run to compute the completeness and contamination parameters of the input genomes.

The input genomes can be annotated with Prokka^26^ or Bakta^27^. As both of these annotators produce results with a similar output structure, it is up to the user to decide which to use for downstream analysis. The default is Prokka, which is also displayed in **Fig. 2**.

Advanced annotation is carried out with the following tools with their briefly stated functions: Interproscan^28^ scans protein signature databases like PFAM, TIGRFAM, and HAMAP. Dbcan^29^ scans carbohydrate active enzymes (cazymes). Eggnog-mapper^30^ provides orthology-based functional annotations. Gapseq^31^ builds gapfilled genome scale metabolic models (GEMs). Antismash^16^ finds biosynthetic gene clusters. Clusterprofiler^32^ computes a pathway enrichment analysis. GTDB-Tk^7^ uses an alignment of ubiquitous proteins to predict species names.

In a clinical setting, the following tools might be useful: Abricate^33^ scans the NCBI^34^, Card^35^, Plasmidfinder^36^, and VFDB^37^ databases for antimicrobial resistance genes and virulence factors. MLST^38,39^ calls multi-locus sequence types, is relevant for an initial grouping when tracking transmission and spread of bacteria.

In terms of phylogenetic analysis: Mashtree^40^, which computes a neighbor-joined tree on the basis of mash distances. Treecluster^41^ which based on customizable presets clusters the mashtree tree. Panaroo^9^ produces a core genome suitable for phylogenetic analysis and defines a pangenome. This core genome is used by the following tools: Fasttree^42^ computes a neighbor joined tree. IQ-TREE^43^ computes a maximum-likelihood tree. Snp-dists^44^ computes the pairwise snp-distances.

The CompareM2 code base with its 1004 source lines of code (SLOC), installation instructions, and documentation are available in a Git repository currently hosted at GitHub: https://github.com/cmkobel/comparem2. The code is published under the GNU Public Licence version 3 which means that anyone who wishes to modify the software can do so if attributing the original authors. If users come up with concrete modifications or extensions, they are welcome to make a pull request on this repository.

### Benchmarking

All running time analyses were run sequentially in random order on an AMD x86-64 “Threadripper Pro” 5995WX 64 cores, 8 memory channels, 512GiB DDR4 3200MHz ECC (8x 64 GiB) and 4 2TB SSDs in raid0. All tests were run with three replicates. Electrical power used consisted of 89% Hydroelectric, 11% wind^45^. The running time was reported by the Snakemake benchmark function. We allowed each pipeline to use a maximum of 32 cores. Statistics and plots were generated using R^46^ v4.3.1 and Tidyverse^18^ v2.0.0. Scripts used to compute the benchmarking results are provided at https://github.com/cmkobel/cm2_benchmark.

### Statistics of analyzed MAGs

Genome sizes were measured with assembly-stats v1.0.1. The MAGs were classified using GTDB v2.3 using database release 214. MAGs from both compared genera (*Methanobrevibacter* and *Prevotella*) were sourced from a project based on ONT R10.4 long read sequencing of the rumen content of 24 male *Bos taurus*. These genomes are accessible here: https://doi.org/10.6084/m9.figshare.26203361

## Declarations

### Ethics approval and consent to participate

Not applicable.

### Consent for publication

Not applicable.

### Availability of data and materials

Bioconda package: https://anaconda.org/bioconda/comparem2 Complete code base repository: https://github.com/cmkobel/comparem2 Documentation: https://comparem2.readthedocs.io Scripts for computing the benchmark results in this paper: https://github.com/cmkobel/ac2_benchmark

### Authors’ contributions

CMK conceptualized, designed, and developed the software. CMK wrote the initial version of the manuscript. VTEA, OØ, NNL, BJW, and PBP assisted in conceptualizing the software and writing the manuscript. All authors read and approved the final manuscript.

## Acknowledgements

We want to thank the initial testers and users of the software for critical feedback: Judith Guitart-Matas, Shashank Gupta, and the 2023 and 2024 student batches of the “BIO326: Genome Sequencing; Tools and Analysis” course at Norwegian University of Life Sciences.

## Author’s information

Carl M. Kobel is a bioinformatician and a PhD candidate in the MEMO group at NMBU, Norway. Carl’s perspective is that microbiomes are largely undervalued and that we should better understand the minute interactions within them. Carl adopts a big data inspired approach, enjoys tinkering with hardware, and building parallelizable bioinformatics pipelines to gain insights into large microbiome datasets.

